# Interrogating the effect of enzyme kinetics on metabolism using differentiable constraint-based models

**DOI:** 10.1101/2022.07.11.499575

**Authors:** St. Elmo Wilken, Mathieu Besançon, Miroslav Kratochvíl, Chilperic Armel Foko Kuate, Christophe Trefois, Wei Gu, Oliver Ebenhöh

## Abstract

Metabolic models are typically characterized by a large number of parameters. Traditionally, metabolic control analysis is applied to differential equation-based models to investigate the sensitivity of predictions to parameters. A corresponding theory for constraint-based models is lacking, due to their formulation as optimization problems. Here, we show that optimal solutions of optimization problems can be efficiently differentiated using constrained optimization duality and implicit differentiation. We use this to calculate the sensitivities of predicted reaction fluxes and enzyme concentrations to turnover numbers in an enzyme-constrained metabolic model of *Escherichia coli*. The sensitivities quantitatively identify rate limiting enzymes and are mathematically precise, unlike current finite difference based approaches used for sensitivity analysis. Further, efficient differentiation of constraint-based models unlocks the ability to use gradient information for parameter estimation. We demonstrate this by improving, genome-wide, the state-of-the-art turnover number estimates for *E. coli*. Finally, we show that this technique can be generalized to arbitrarily complex models. By differentiating the optimal solution of a model incorporating both thermodynamic and kinetic rate equations, the effect of metabolite concentrations on biomass growth can be elucidated. We benchmark these metabolite sensitivities against a large experimental gene knockdown study, and find good alignment between the predicted sensitivities and *in vivo* metabolome changes. In sum, we demonstrate several applications of differentiating optimal solutions of constraint-based metabolic models, and show how it connects to classic metabolic control analysis.

## 2 Introduction

Biological processes are typically characterized by a large number of parameters. In the context of constraint-based models, these can include enzyme kinetics and thermodynamic constants. Databases of *in vitro* measurements [1, 2] organize and expose decades of research, allowing these parameters to be included in modern models [3, 4, 5]. Incorporating parameters into constraint-based models has been shown to increase their predictive capabilities [6, 7, 8, 9]. For example, classic flux balance analysis is incapable of modeling overflow metabolism without ad hoc assumptions [10], but if enzyme turnover numbers and capacity limitations are incorporated, this phenomenon can be mechanistically modeled [11].

Recently, *in vivo* estimates of enzyme turnover numbers have become available through integrating omics measurements with models [12]. These estimates substantially increase the quantitative accuracy of predictions afforded by enzyme-constrained metabolic models [13]. Machine learning techniques have also made use of these omics driven measurements to extend kinetic parameter estimates, including turnover numbers and Michaelis constants, to the genome-scale [14, 15]. With these advancements it is becoming feasible to construct increasingly detailed metabolic models [16, 17]. However, an unaddressed question is the sensitivity of model predictions to parameters.

Metabolic control analysis (MCA) encompasses a rich theory for estimating parameter sensitivities in ordinary differential equation (ODE) based models [18]. However, a corresponding theory for constraint-based models is lacking. In the latter case, a finite difference based procedure is typically used to estimate the effect of a perturbed parameter on an optimal solution relative to a reference solution [19, 20, 21]. Consequently, for each parameter, a new optimization problem needs to be solved. This suggests that the time required to map the complete sensitivity grows with with the product of the optimizer solve time and the number of parameters. In contrast, MCA implicitly differentiates the ODE-based model, yielding all the sensitivities directly after solving a single system of linear algebraic equations.

Here, we show that an optimal solution of a constraint-based model can be likewise implicitly differentiated, and demonstrate several applications. Calculating the so-called flux control coefficients [19] simplifies to finding the derivatives of the optimization variables to parameters, analogously to classic MCA. Moreover, the ability to differentiate a constraint-based metabolic model allows it to be efficiently embedded in more complex optimization schemes. We use this to solve a bilevel optimization problem, which seeks to fit apparent turnover numbers to an enzyme-constrained model by minimizing the difference between model predictions and observations. Finally, we show that the method is easily extendable to problems of arbitrary complexity by comparing the predicted sensitivity of intracellular metabolite concentrations to biomass growth in a model that incorporates both thermodynamic constraints, as well as Michaelis-Menten reaction kinetics.

## 3 Results

A brief note on nomenclature: one dimensional variables are denoted like *x*, vectors like **x**, and matrices like **X**. Further, qualitative models are shown in Section 3, but the full models are shown in Section 5.

### 3.1 Extending metabolic control analysis to constraint-based models

Metabolic control analysis (MCA) is a quantitative technique widely used to gauge the sensitivity of variables to parameters in dynamical systems [22, 18]. Briefly, suppose a biochemical process is governed by a system of differential equations,

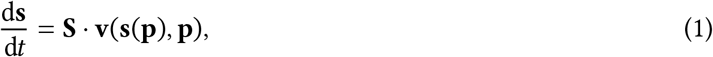

which relate reaction fluxes (**v**) to metabolite concentrations (**s**) through parameters (**p**), using the stoichiometric matrix (**S**). We can define the function **f**_SS_,

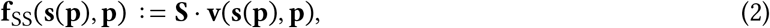

and at steady state,

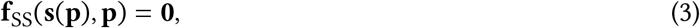

which implies an implicit relationship between **p**, and **s**. Consequently, implicit differentiation of Equation (3) yields,

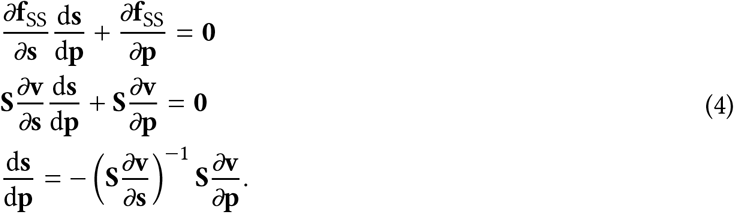

Thus, 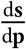 represents the sensitivity of the variables to parameters at steady state, and from here the classic flux and concentration control coefficients, as well as elasticities, can be calculated [23]. Taken together, these sensitivities are calculated by applying the implicit function theorem to a steady state solution of some governing differential equation. Moreover, if the derivatives of the constituent functions are known, then finding the sensitivities only require the solution of a single system of linear algebraic equations.

#### Constraint-based metabolic control analysis

Typically, constraint-based models without integer variables can be expressed as either convex linear or convex quadratic programs [24], which have the general form,

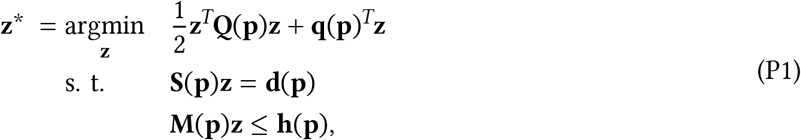

where **Q** is a positive semidefinite matrix. Here, **z**^∗^ represents an optimum of the variables (often fluxes) given parameters (**p**), which can include enzyme turnover numbers, Michaelis constants, etc. The dependence of the optimization program on parameters is usually not shown, but here it has been made explicit. Normally, **S** represents the stoichiometric matrix, **d** = **0**, and **M, h** are bounds placed on the variables to enforce physiological constraints (like reaction directions, flux bounds, etc.). For convenience, define,

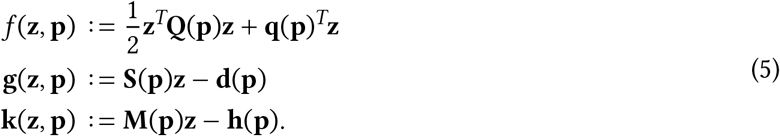

Optimization theory holds that for a convex problem, like Problem (P1), the stationarity, primal feasibility and complementary slackness conditions (first-order Karush-Kuhn-Tucker conditions), shown in Equation (6), are both necessary and sufficient for optimality [25],

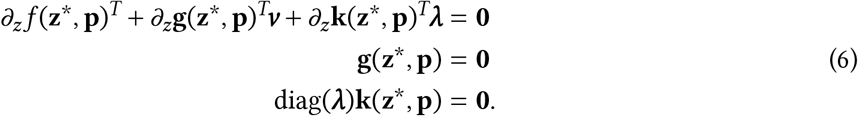

Here, *v* and *λ* are the equality and inequality dual variables (generalizations of Lagrange multipliers), and diag(⋅) denotes the diagonal matrix formed from a vector. To highlight the implicit dependence of the optimum variables on the parameters, one may rewrite the optimality conditions by substituting the definitions of Equation (5) into Equation (6), and define a new function, **f**, which resembles **f**_SS_,

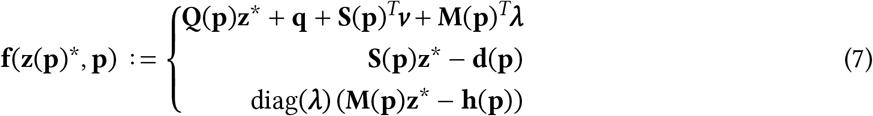

Thus, at the optimum,

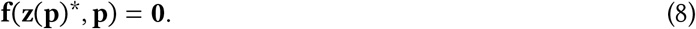

This suggests that implicitly differentiating **f** will yield the derivatives of the optimization variables to their parameters [26, 27, 28, 29]. Specifically, the similarity between **f**_SS_ in Equation (3) and **f** in Equation (8), reveals the connection point between classic MCA and its constraint-based model counterpart. By implicit differentiation of the optimality condition function,

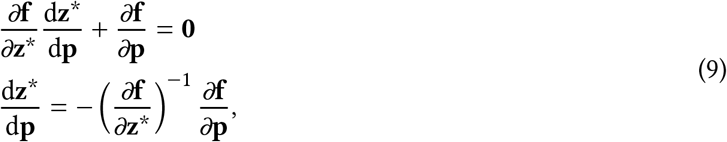

the sensitivity of the variables at the optimum to model parameters, 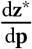, can be found. Moreover, similar to the classic MCA approach, if the derivatives of the constituent functions are known, then the constraint-based metabolic control analysis (CB-MCA) procedure is computationally efficient, because only a single linear system of equations needs to be solved (subsequent to the optimization) to calculate *all* the derivatives. Encouragingly, recent developments in automatic and symbolic differentiation allows these constituent function derivatives to be calculated efficiently for arbitrary problems automatically [30, 31, 32].

### 3.2 Metabolic control analysis of an enzyme-constrained metabolic model

Constraint-based models represent a scalable framework to interrogate microbial metabolism. Incorporating enzyme capacity and rate limitations into these models unlocks the ability to mechanistically explain resource allocation phenomena, including overflow metabolism [11]. Various frameworks have been proposed to model enzyme limitations [4, 6, 33]. Qualitatively, Problem (P2) summarizes their main features,

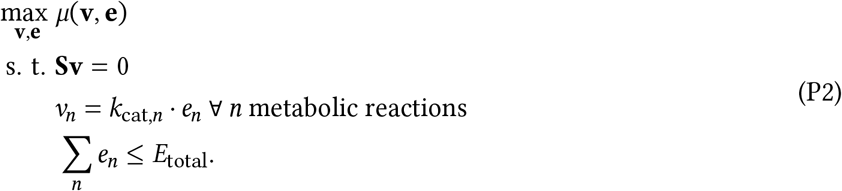

Importantly, these models hold that the flux, *ν*, through each reaction is proportional to the enzyme concentration, *e*, that catalyzes the reaction, and its associated turnover number, *k*_cat_. Additionally, the total proteome capacity is limited by *E*_total_, and *µ* represents the biomass objective function. Typically, the turnover numbers are taken as constants, often inferred from databases [21] or estimated [12, 14]. With the CB-MCA framework introduced earlier, it is possible to efficiently estimate the sensitivity of the predicted intracellular fluxes and enzyme concentrations to the turnover numbers.

Figure 1 shows the predicted reaction fluxes and enzyme concentrations of *Escherichia coli* growing aerobically on glucose in minimal media using the GECKO formulation [6] of Problem (P2), together with turnover numbers for all the metabolic enzymes, generated using a machine learning based approach [34].

**Figure 1:**
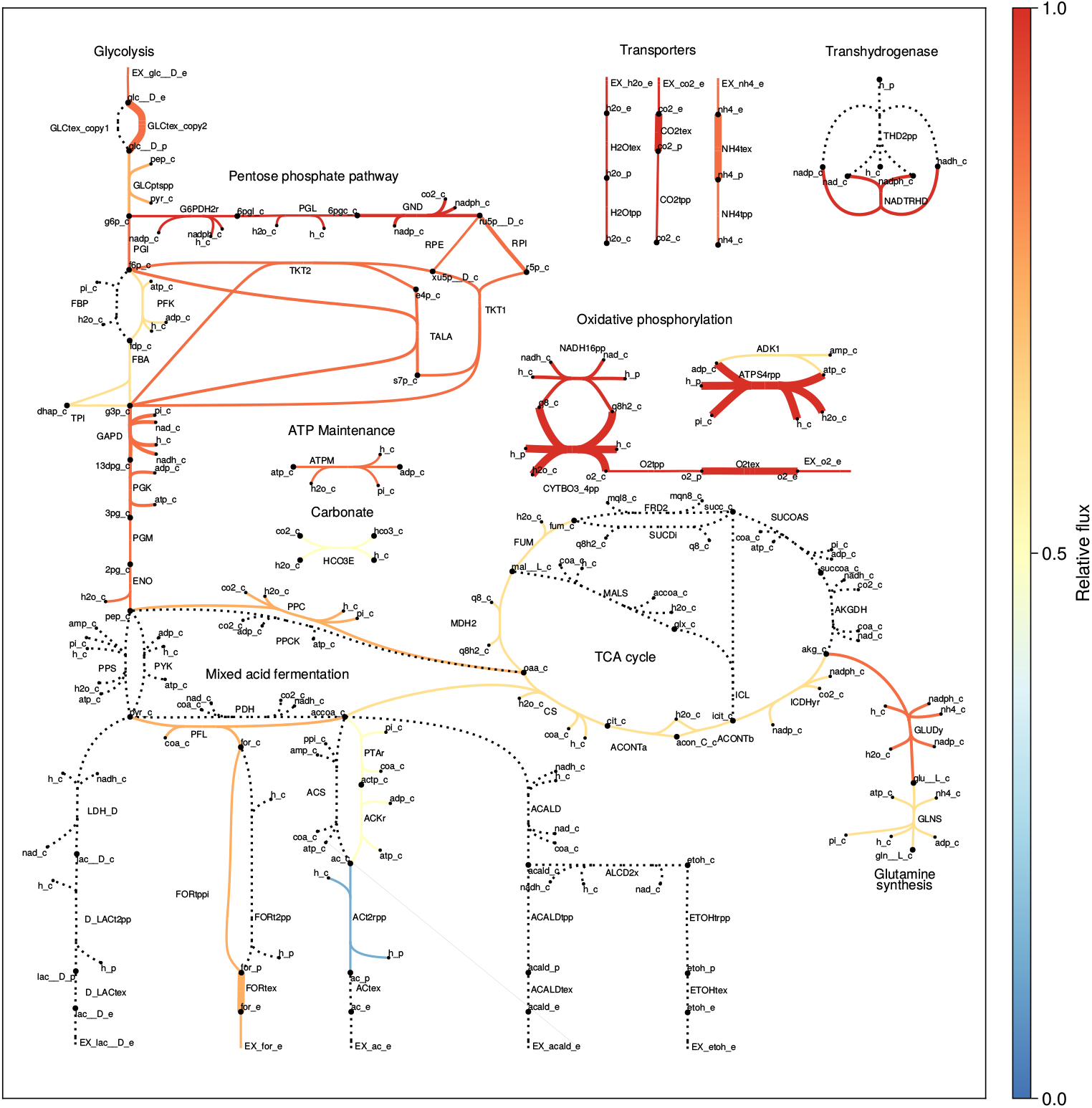
Enzyme-constrained metabolic models can predict intracellular fluxes and protein concentrations. The core carbon metabolism of *E. coli* growing aerobically on glucose in minimal media is shown here. Enzyme limitations were incorporated into its latest metabolic model [35] using the GECKO formulation [6], and turnover numbers were taken from a genome-wide dataset [34]. Reaction fluxes are colored according to their relative absolute magnitude, and reaction edge width is scaled relative to the predicted enzyme concentration. Metabolically inactive reactions are dashed. This metabolic realization (fluxes and enzyme concentrations) is differentiated in Figure 2.

The sensitivities of both the reaction fluxes (Figure 2 A), as well as the enzyme concentrations (Figure 2 B), to the turnover numbers can be found by differentiating the optimal solution, shown in Figure 1. To prevent differentiability issues caused by the potential non-invertibility of 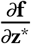, only active reactions (those that are not dashed in Figure 1) and expressed enzymes are differentiated (see Section 4 for more details).

**Figure 2:**
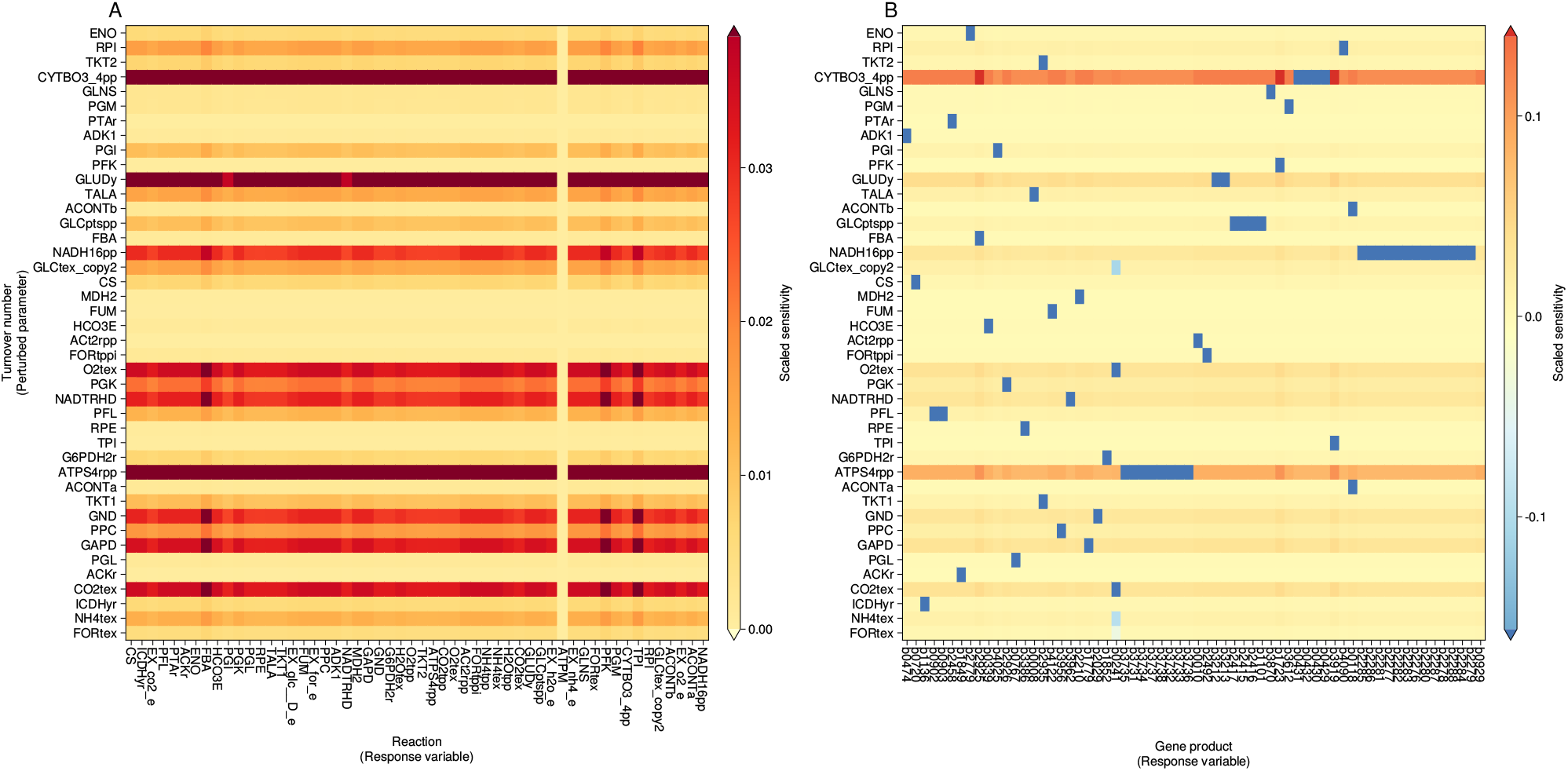
Scaled sensitivities of predicted reaction fluxes and gene product concentrations to enzyme turnover numbers. *E. coli* model iML1515 was simulated under aerobic, glucose fed conditions, and subsequently the active solution, shown in Figure 1, was differentiated to yield these sensitivities. The sensitivity of variable *z*_*i*_ to parameter *p*_*j*_ shown here are scaled, 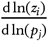. For simplicity, only enzymes in the core carbon metabolism are shown here.

In Figure 2 A, the flux sensitivities indicate that cytochrome oxidase (*CYTBO3_4pp*), glutamate dehydrogenase (*GLUDy*), and ATP synthase (*ATPS4rpp*) exert the most control over the optimal solution. Quantitatively, increasing their associated turnover numbers will increase flux across all active reactions, except the maintenance reaction (*ATPM*). Figure 2 B, which shows the sensitivities of the predicted gene product concentrations to the turnover numbers, illustrates why this is expected. Taking cytochrome oxidase as an example, the sensitivities of its subunits to its turnover number are negative, while all other gene products have a positive sensitivity to the turnover number of cytochrome oxidase. If the turnover number for cytochrome oxidase increases, less enzyme is required to catalyze the same amount of flux, consequently, more enzyme can be distributed to other reactions while satisfying the enzyme capacity bound. Consequently, the flux through metabolism increases. In contrast, the maintenance reaction is an ATP sink, and increasing flux through it can only decrease the objective function. Hence, when biomass is maximized it is forced to its lower bound, and thus its sensitivity is zero to all turnover numbers. Interestingly, comparing the model-based flux sensitivity of an enzyme to its turnover number against the measured absolute protein abundance of the respective enzyme under the same conditions, we find a positive linear relationship (*R*^2^ = 0.49, see Figure S1). This suggests that the control coefficients accurately identify reactions that exert control over fluxes in metabolism.

Beyond identifying flux controlling enzymes, the sensitivities shown in Figure 2 also indicate the relative flux increase a reaction would experience if the associated turnover number were to be increased. However, unlike classic MCA, the summation theorem does not have a clear analog in the constraint-based case. Despite this difference, the sensitivities shown here are *exactly* the same as the finite difference based sensitivities, called flux control coefficients, introduced in other work [19, 21] (see Figure S2). Finally, inspecting all the sensitivities (see Figure S3) shows that most parameters have a small scaled flux sensitivity (≤ 10^−2^), suggesting that precise turnover numbers for all metabolic enzymes are likely not critically important.

### 3.3 Differentiating an enzyme-constrained metabolic model to fit turnover numbers to measured data

Incorporating enzyme turnover numbers and capacity limitations into flux constraints has been shown to dramatically increase the predictive capabilities of constraint-based models [13]. Unfortunately, it is experimentally infeasible to clone and express each enzyme a cell possesses for individual characterization, in part due to the large number of protein coding genes contained in even minimal-sized microbes. Recent approaches to tackle this characterization problem for enzyme turnover numbers include machine learning [14, 34], and constraint-based metabolic modeling [12] techniques. The former method makes use of a variety of enzyme specific training data (crucially, including measured kinetic data), and is capable of predicting enzyme turnover numbers at the genome-scale for a specific organism, albeit through a black-box technique. The latter method couples omics measurements to a mechanistic model, and can directly estimate the apparent turnover numbers of expressed enzymes. Despite the usefulness of these *in vivo* estimated turnover numbers, only *Saccharomyces cerevisiae* [13] and *Escherichia coli* [14] have been subject to such genome-scale based approaches.

In the constraint-based model approach to estimate apparent turnover numbers, flux balance analysis (FBA) *without enzyme constraints* is used to estimate the internal reaction fluxes. Measured fluxes are used to constrain the model to increase the accuracy of the predicted fluxes. Apparent turnover numbers are then estimated by,

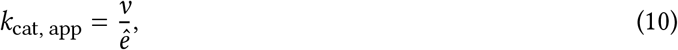

where *ê* is the intracellular measured protein concentration of the enzyme catalyzing the relevant reaction. By sampling many metabolic conditions, the true turnover number can be estimated by taking the maximum of all the apparent turnover numbers. The primary drawback of this approach is that quantitative proteomic measurements do not span the entire metabolism, limiting which reaction turnover numbers (typically homomeric enzymes) can be estimated in this way.

Fundamentally, conventional FBA is used instead of an enzyme-constrained variant, because once enzyme kinetics, intracellular fluxes, and enzyme concentrations are incorporated into a model as variables, it becomes a nonlinear optimization problem, which is challenging to solve at the genome-scale. A practical workaround for this issue is to set the kinetic constants, which need to be estimated, as parameters, and only optimize over the fluxes and enzyme concentrations. For a given set of turnover numbers, **k**_cat_, the mean squared relative error between the model predicted intracellular fluxes and enzyme concentrations, and their measured counterparts, can be minimized, as shown qualitatively in Problem (P3),

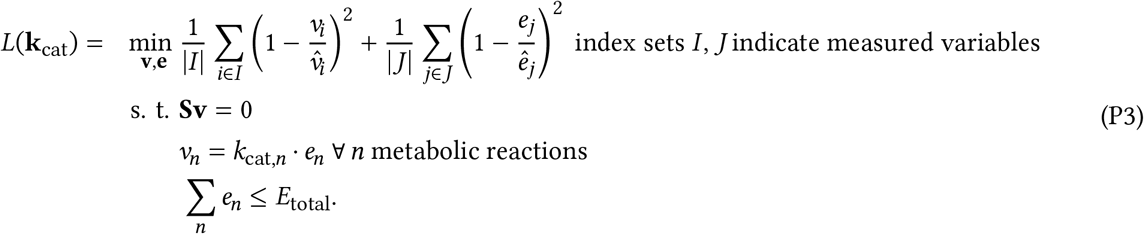

The benefit of this approach is that both missing fluxes *and* missing enzyme concentrations can be imputed by solving the model. This broadens the scope of apparent turnover numbers that can be estimated by using the model-based approach. Additionally, this optimization problem is a convex quadratic program with only diagonal Hessian terms, for which highly efficient methods and corresponding solvers exist [36]. All that remains is to change **k**_cat_ to minimize the error function, *L*, in Problem (P3), to find the turnover numbers that best fit the measured data.

A computationally tractable approach to reduce the error induced by inaccurate turnover number estimates is to perform gradient descent on *L*, which requires that its derivatives can be efficiently calculated. Thus, by using the differentiation techniques introduced earlier, the bilevel optimization problem of finding the set of turnover numbers minimizing *L* subject to Problem (P3) can be solved. The optimum corresponds to the turnover numbers that minimize the discrepancy between model predictions and measured data.

Using measured intracellular fluxes and protein concentrations of *E. coli* under various culture conditions [34], this technique can be tested. In Panel A of Figure 3, the mean squared relative error, *L*, is shown over multiple gradient descent iterations. Starting from the turnover numbers estimated by the machine learning approach, the algorithm successfully changes them to reduce the error. Panel B shows the resultant turnover numbers for enzymes in glycolysis, and Panel C their derivatives. We observe that both derivatives and errors became smaller over iterations, indicating that the turnover numbers converge.

**Figure 3:**
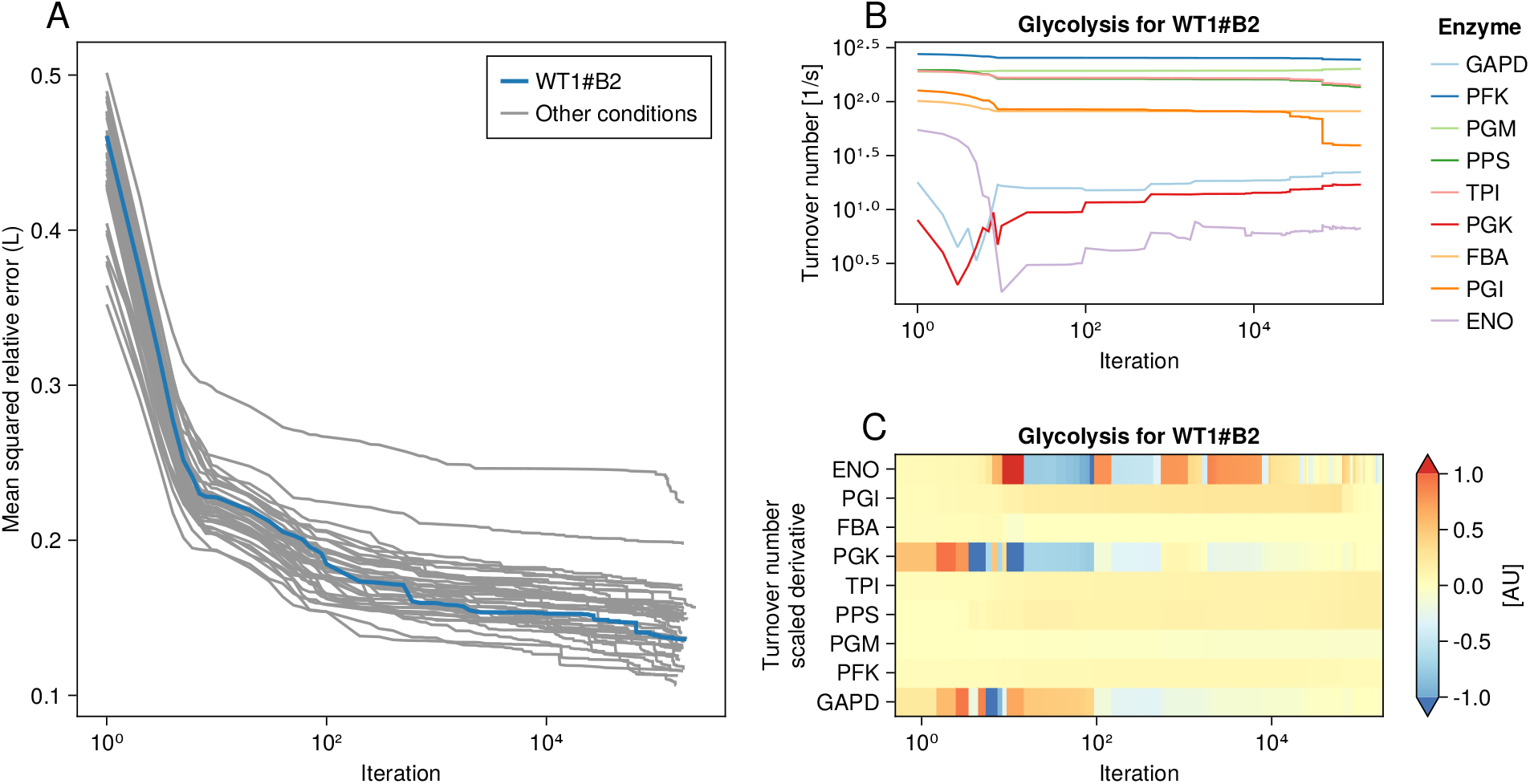
Turnover number estimates can be improved by minimizing the difference between observations and model predictions. Panel A highlights the mean squared relative error of a specific culture condition in a published dataset (WT1#B2: wild type cell, condition replicate 1, technical replicate 2), the other conditions are shown in grey. Panel B shows how the turnover number estimates in glycolysis changes over gradient descent iterations, subject to the derivatives in Panel C.

In keeping with previous work [12], the maximum turnover number estimate from all the culture conditions descended over in Figure 3 was taken as the best estimate of the true enzyme turnover number. Panel A in Figure 4 shows that using these improved turnover number estimates, compared to the state-of-the-art machine learning generated estimates, increases the accuracy of model predictions against experimental measurements by 30 ± 3%. Comparing the improved turnover number estimates to a subset of those found in BRENDA [12], the coefficient of determination (*R*^2^) is 0.39, suggesting that they are in line with the *in vitro* data. The final set of improved turnover number estimates is listed in Supplementary Dataset 1.

**Figure 4:**
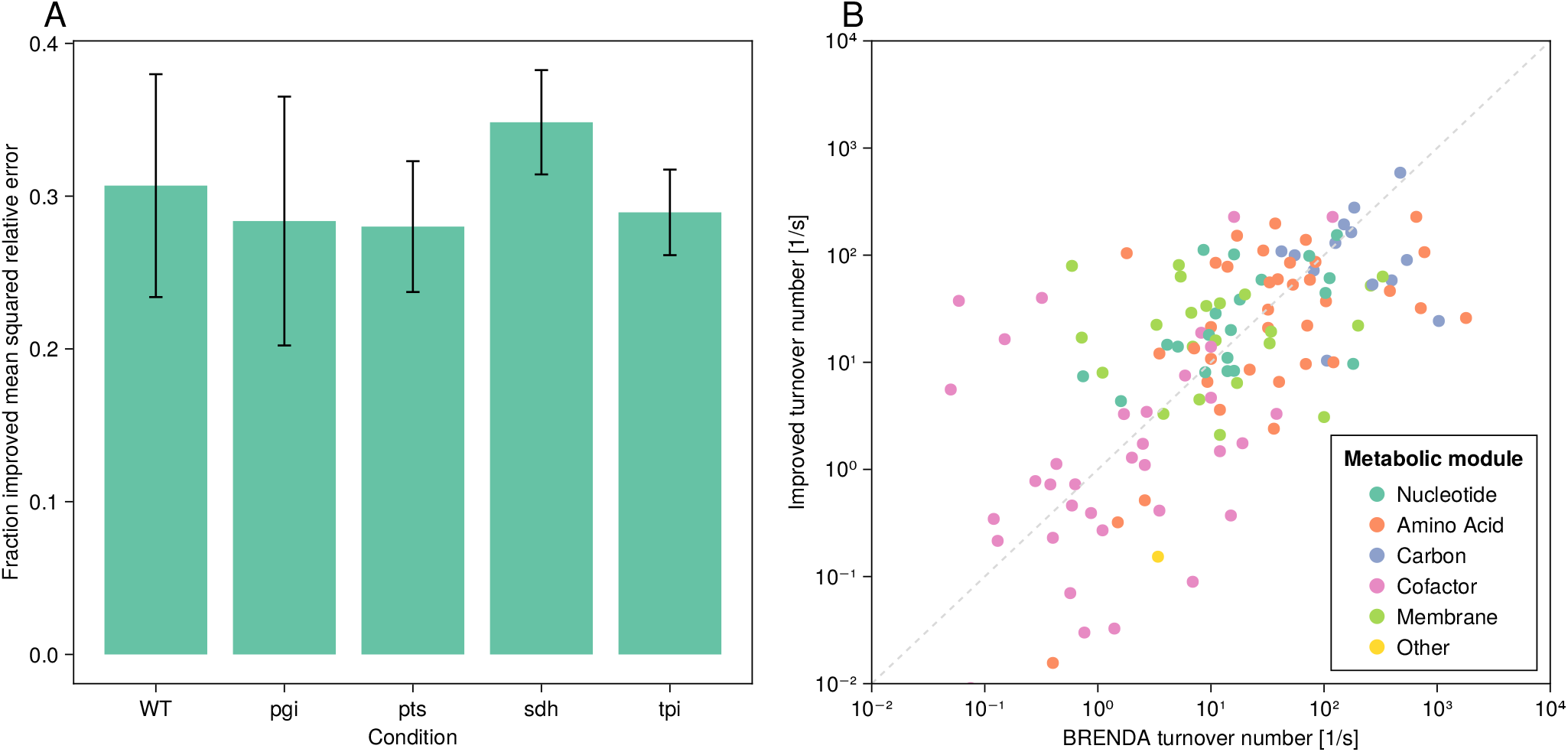
Improving the machine learning generated turnover number estimates with the differentiable modeling approach causes the accuracy of enzyme-constrained model predictions to increase. Panel A shows the fractional increase in accuracy achieved when using the improved turnover number estimates compared to their state-of-the-art machine learning generated counterparts to predict intracellular fluxes and enzyme concentrations using the latest *E. coli* metabolic model under various genetic knockout conditions (WT: wild type, pgi: glucose-6-phosphate isomerase, pts: glucose phosphotransferase system, sdh: succinate dehydrogenase, tpi: triose-phosphate isomerase). To make the comparison fair, for each test condition its data is left out of the dataset used by the gradient descent technique (hold-out method). However, the published machine learning data is used as is, which incorporates all the available data, making the test more conservative against the gradient descent technique. Error bars reflect the standard deviation of the error function, *L*, for all the experimental replicates in each condition. Panel B shows a comparison of the improved turnover numbers to those found in BRENDA for a curated list of enzymes [12], grouped by the metabolic module they occur in.

### 3.4 Differentiable metabolic models can account for enzyme saturation and thermodynamic effects

While the aforementioned enzyme constraints represent physiologically realistic simplifications, enzyme kinetics are typically much more complex *in vivo*. Regulatory, thermodynamic, and saturation effects can dramatically alter the maximum rate at which an enzyme can metabolize its substrate [37]. In particular, metabolite concentrations directly affect the thermodynamic driving force, and enzyme active site occupancy [38]. Recently, CRISPRi was used to knockdown gene expression to reduce the concentration of various enzymes in *E. coli* with the goal of understanding how the metabolome responds to enzyme limitations. The experimental data showed that the substrate metabolites of the enzyme that was throttled tended to increase in concentration, unless another form of regulation (e.g. allosteric, etc.) was available [39]. Thus, by altering its metabolome, *E. coli* compensated for the knockdowns, and the deleterious effect on growth rate was reduced.

Incorporating metabolite concentrations as variables into constraint-based models is challenging, because thermodynamic and saturation effects are inherently nonlinear [40]. Typically, either fluxes (e.g., in max-min driving force analysis [3]) or metabolite concentrations (e.g., in flux balance analysis [24]) are abstracted away into the model. This approach precludes the ability to investigate the sensitivity of metabolite concentration on flux and enzyme concentration predictions. To address this, we extend the simple enzyme-constrained model given in Problem (P2) to include metabolite concentrations as parameters, as shown in Problem (P4),

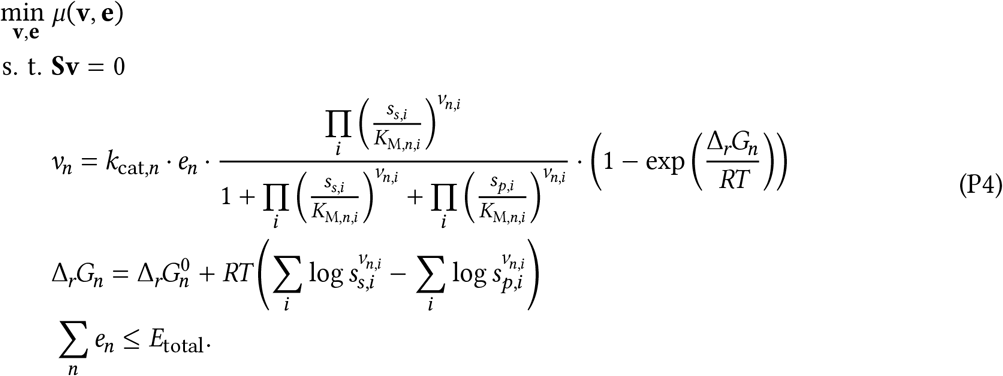

Here, *K*_M,*n,i*_ and *v*_*n,i*_ denote the Michaelis constant and the stoichiometric coefficient of reaction *n* and metabolite *i*, respectively. Further, *s*_*s,i*_ or *s*_*p,i*_ indicate the substrate or product concentration of metabolite *i* relative to the associated reaction, and 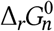 is the standard Gibbs free energy change of reaction *n* [37]. This model accounts for both saturation and thermodynamic effects, but ignores regulation. Since the metabolite concentrations are taken as parameters (estimated as described in Section 5), the problem remains convex, and is computationally tractable.

Through the addition of Michaelis-Menten like kinetics in Problem (P4), we hypothesize that the experimentally observed metabolome changes can be understood by investigating the sensitivity of the biomass function to intracellular metabolite concentrations under knockdown conditions. Specifically, metabolites with high sensitivity are likely to exert larger control on the biomass function, suggesting that they could be used to counteract the effect of the knockdowns. Thus, we test if the model can recapitulate the observed trend that substrates to the throttled enzyme increase in concentration, by evaluating if the substrate metabolites have high sensitivity.

Figure 5 shows these sensitivities for a representative selection of knockdown conditions (all other conditions are shown in Figure S4, but are broadly similar to the ones shown here). Each simulated knockdown condition constrains the corresponding gene concentration to be five-fold less than it would be under wild-type conditions. Subsequently, the model in Problem (P4) is simulated, and differentiated. The resulting sensitivities of the biomass function to intracellular metabolites are compared to experimentally measured metabolite concentration fold changes subject to the same knockdowns [39].

**Figure 5:**
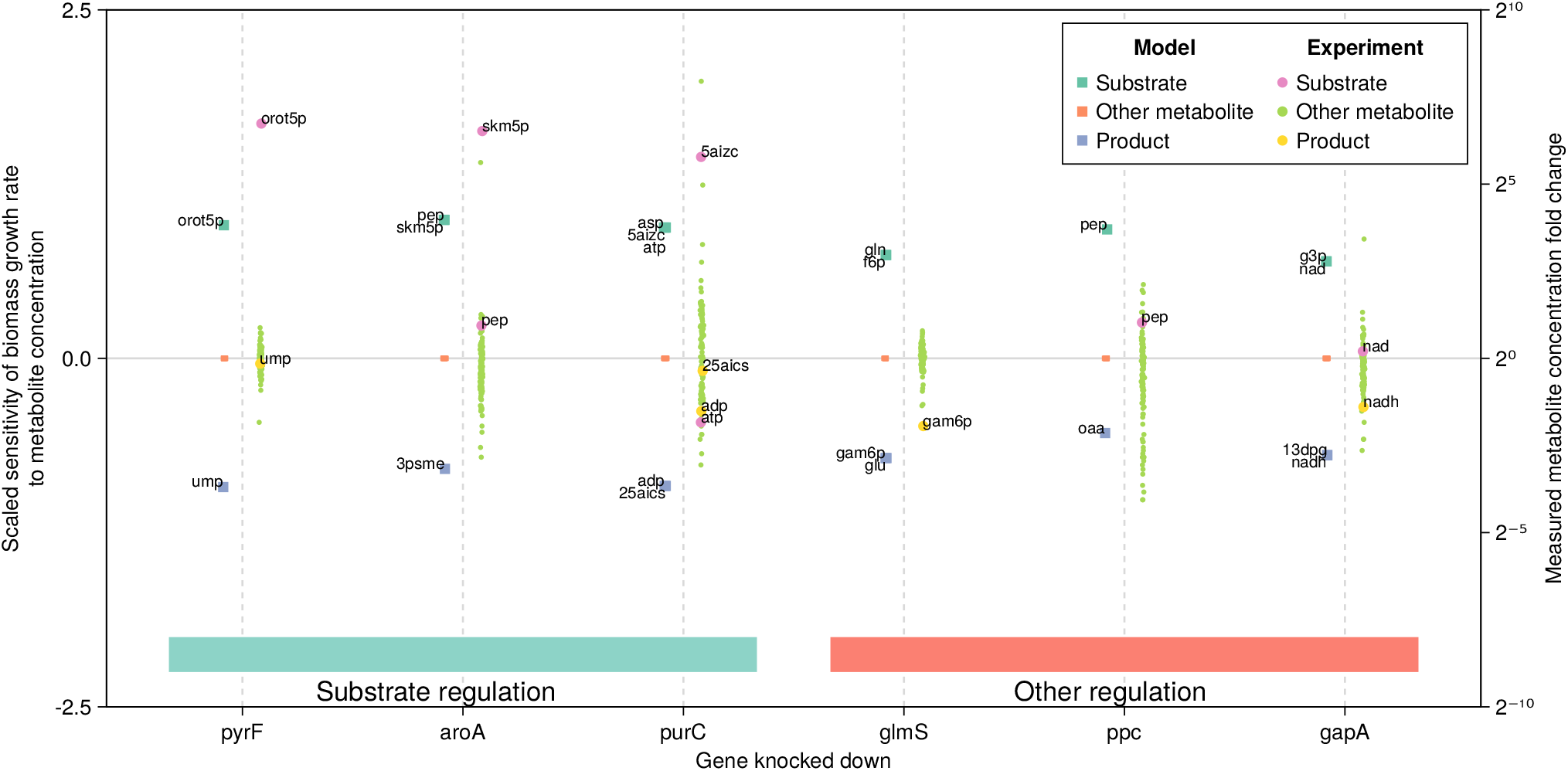
Predicted biomass sensitivity to metabolites aligns with experimentally measured metabolome changes under genetic perturbations. Each gene knockdown has two datasets associated with it, separated by a vertical dotted line. On the left y-axis, the simulated sensitivities of the biomass function to metabolite concentrations is shown after the associated gene is constrained to be five-fold less abundant than the wild type system. Substrate and product metabolite sensitivities are highlighted. On the right y-axis, experimental data of intracellular metabolite concentration fold changes observed after CRISPRi knockdowns of the associated genes are shown [39]. Substrate and product metabolites are also highlighted if measured. The gene abundance constraints experimentally observed match those simulated *in silico*. The groupings denoted by the horizontal blocks near the bottom of the figure separate the primary compensating mechanism observed *in vivo*. Briefly, “‘Substrate regulation” denotes enzymes where the measured substrate metabolites played a large role in controlling the kinetics. “Other regulation” denotes cases where substrate concentration played a smaller role, e.g. allosteric regulation compensated for the knockdown, etc. Each gene catalyzes the following reactions: Orotidine-5’-phosphate decarboxylase (*pyrF*), 3-phosphoshikimate 1-carboxyvinyltransferase (*aroA*), Phosphoribosylaminoimidazolesuccinocarboxamide synthase (*purC*), Glutamine-fructose-6-phosphate transaminase (*glmS*), Phosphoenolpyruvate carboxylase (*ppc*), Glyceraldehyde-3-phosphate dehydrogenase (*gapA*).

The simulated sensitivity results, shown in Figure 5, indicate that the substrate and product metabolites of the throttled enzyme have the largest sensitivities. The sensitivity of most other intracellular metabolites is small (≤ 10^−6^). This suggests that by increasing the substrate concentration (positive sensitivity), or decreasing the product concentration (negative sensitivity), of the throttled enzyme, the flux through the biomass reaction can be increased. Thus, the model recapitulates the experimental observations, although it is agnostic between product or substrate metabolites. Specifically, for enzymes where reagent metabolites compensate for enzyme limitations (e.g., *pyrF, aroA, purC*), the sign of the sensitivities aligns with the observed metabolome changes.

However, since Problem (P4) does not include regulatory mechanisms, it is incapable of accounting for allosteric control, as observed in *glmS* and *ppc* (allosterically regulated through glucosamine-phosphate, and aspartate and malate, respectively) [39]. Instead, the model predicts that the substrate and product metabolites would compensate for the knockdown. Similarly, for *gapA*, where the compensation mechanism is unclear [39], the model predicts saturation compensation, because this is the only explanatory mechanism built into the mathematical structure of the model.

## 4 Discussion

### Constraint-based metabolic control analysis

Classic metabolic control analysis provides a precise description of the sensitivities of variables to parameters in ODE-based models. However, these models are exceedingly challenging to parameterize at the genome-scale, limiting their applicability to smaller systems [41]. Constraint-based models make use of substantially less parameters, which widens their scope. However, they are cast as optimization problems, superficially occluding the direct application of MCA to them. Here, we show that precisely the same mathematical technique, as is used in ODE-based models, can be used calculate the sensitivities of variables in constraint-based models to their parameters. This dramatically improves upon the finite difference based approach currently used (a single parameter is perturbed and the difference between a reference and response solution is used to estimate sensitivities). Moreover, by making the definition of the derivatives precise, metabolic degeneracy must be dealt with explicitly, clarifying the meaning of the resultant sensitivities.

In particular, models making use of flux balance analysis, including their enzyme-constrained variants, invariably need to sample different solutions to form a complete picture of the metabolic potential of an organism [42]. Fundamentally, this is because they are typically solved through linear programming, which only guarantees a unique objective is attained. This presents problems for finding meaningful derivatives, as the variables used in the derivative calculations are not guaranteed to be unique. In the finite difference based approach, this effect is ignored. However, the implicit differentiation technique introduced here reveals that the Jacobian, 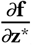, is singular in these cases, because there are linearly dependent (degenerate) metabolic states. Unless special steps are taken to ensure a unique basis forms the optimum objective, the Jacobian will not be invertible, and thus the sensitivities are not well defined. In general, a small quadratic regularizing term can be added to the linear program, turning it into a quadratic program, with uniqueness guarantees on both the objective and variables. While this transforms it into a strongly convex program that can be differentiated [43], the cost is solving a more complicated problem. Alternatively, by solving the model and removing metabolically inactive variables (i.e. removing the metabolic degeneracy), a unique optimum basis can be found [44]. Subsequently, the Jacobian is full rank, and the solution may be differentiated. In sum, either procedure results in well defined derivatives, making their subsequent interpretation simpler (see Section S1 for more details).

In light of the foregoing, different metabolic realizations (satisfying the same optimality condition) can yield different sensitivities (demonstrated in Figure S5). While most reactions display small variations in sensitivity, some metabolic modules, including the tricarboxylic acid cycle and nucleotide metabolism, can be substantially different. This highlights the importance of preferring the implicit differentiation approach: it ensures that the resulting sensitivities are well-defined by forcing the reference solution to be defined prior to differentiation. Finally, the clear connection to classic MCA and the associated derivatives suggests deeper connections between theoretical advances and specific models are possible. For example, marginal fitness cost, the derivative of protein cost to metabolite concentration as introduced in growth balance analysis [45], can be simply extended to arbitrary metabolic models within this framework, as the necessary derivatives can now be calculated automatically.

### Estimating enzyme turnover numbers

Enzyme kinetic databases collect data gathered over decades and can be used for model parameterization [2, 46]. However, these *in vitro* estimates are typically noisy [47] and *in vivo* generated data tends to generate better predictions when used in models [14, 13]. Despite the clear benefits of using *in vivo* data to estimate enzyme kinetics, challenges remain in generating accurate parameter estimates across all metabolically active enzymes.

Conceptually, it is possible to *only* use measured intracellular fluxes and protein abundances to estimate turnover numbers. However, the primary drawback of this approach is that proteomic and intracellular flux measurements do not span the entire metabolism. Thus, measurement gaps need to be filled, motivating the use of a model. In previous work, FBA was used to find missing fluxes, with the shortcoming that missing enzyme concentrations could not be imputed [12]. Subsequently, machine learning was used to estimate the remaining turnover numbers. In this work, the differentiability of an enzyme-constrained model was leveraged to refine the machine learning estimates (which were used as the initial conditions for the gradient descent algorithm). Measurement gaps were imputed by the model for all metabolically active enzymes. A caveat to this approach is that the bilevel optimization problem being solved is nonconvex, suggesting multiple solutions may exist. Different initial conditions may give rise to different sets of improved turnover number estimates. While randomly sampled starting points also converged to low errors (see Figure S6), using the machine learning estimated turnover numbers as starting points tended to yield the lowest error over the gradient descent iterations. This suggests that the machine learning approach generated good initial estimates. Comparing the final improved turnover number estimates to their machine learning counterparts reveals that the former are higher on average (see Figure S7). Since the improved turnover numbers increased the predictive accuracy of the model, it indicates that the machine learning approach underestimated the true turnover numbers.

The bilevel algorithm introduced here imputes missing enzyme concentrations while descending on the associated turnover numbers. Consequently, there is no clear upper bound for the associated turnover numbers of the missing enzymes. Thus, it is possible for the algorithm to increase the turnover numbers of these enzymes to physiologically unrealistic values to minimize the amount of enzyme required to catalyze the predicted fluxes. Practically, this is not too concerning because the objective function of the inner problem merely seeks to fit data, and not e.g., maximize growth (the latter objective could more easily lead to unrealistically large turnover number estimates). Moreover, the gradient descent algorithm changes the turnover numbers of all the enzymes simultaneously, thus the imputed enzyme concentrations effectively get changed relative to the initial estimates, attenuating dramatic changes in turnover number estimates of single reactions. Thus, given the increasing number of machine learning generated turnover number estimates [14, 48], this algorithm is well suited as a refinement step to increase their accuracy.

While data-intensive, this approach promises to unlock the ability to rapidly estimate enzyme kinetics at the genome-scale for any organism in a relatively short time. This addresses the pressing need for kinetic parameters spanning a wider selection of organisms [49]. The model based technique has the advantage of being a transparent process, making the fitting procedure conceptually easier to understand, although it cannot estimate turnover numbers of enzymes that are not expressed, unlike the machine learning-based approach. Consequently, only 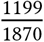 of the metabolic reactions were assigned turnover number estimates using the gradient descent-based approach (see Supplementary Dataset 1).

### Sensitivity analysis of a complex model

Given the broad applicability of differentiable constraint-based models, we demonstrate that calculating their sensitivities is straightforward, even when incorporating thermodynamic and kinetic constraints. Such a model was used to determine the sensitivities of intracellular metabolite concentrations to biomass growth. This model was partially able to recapitulate measured trends in metabolite changes after genetic perturbations, likely hampered by regulatory details that were not modeled. Indeed, the model was incapable of predicting allosteric regulation, because the kinetic rate equations did not incorporate this phenomenon. However, saturation effects were captured by this model, suggesting that the approach taken here revealed physiologically realistic control levers cells use to modulate intracellular fluxes.

In light of the procedure used to estimate enzyme turnover numbers *in vivo*, the foregoing results raise the question: can a similar gradient descent based technique be used to find Michaelis constants or standard Gibbs free energy of reactions at the genome-scale? Recent strides in metabolomics allow broad intracellular metabolite concentrations to be measured in addition to fluxes and protein abundances [50, 51]. This suggests that a similar model-based fitting procedure could be used to estimate kinetic parameters beyond just turnover numbers. A potential stumbling block for this would be the nonlinearity introduced by setting metabolite concentrations as variables (opposed to parameters as used in this work), which could make optimization challenging. Nevertheless, the ability to efficiently differentiate the resultant solution could be leveraged to estimate elusive kinetic parameters.

In sum, differentiable constraint-based models provide a systematic basis to explore the effect parameters have on metabolic phenotypes in constraint-based models. We have shown the link between classic MCA, and its constraint-based counterpart, CB-MCA. Further, we demonstrate several applications where the differentiability of optimization problems can help refine parameter estimates, or elucidate the response of their perturbation on models. Increasingly complex metabolic models are being developed, which usually depend on a wide variety of parameters. Sensitivity analysis could help justify the inclusion of theoretical mechanisms in these models, accelerating the pace of systematically understanding cellular phenomena.

## 5 Materials and Methods

All software and worflows used to generate the results in this work can be accessed here https://gitlab.com/qtb-hhu/differentiablemetabolismcode. In this section a high level summary is presented; see the readme in the software repository for more detailed information.

### 5.1 Model construction, parameter sources, and software used

The latest *E. coli* metabolic model, iML1515 [35], was used in all simulations. The core biomass function was optimized, unless otherwise noted (e.g. for the gradient descent in Section 3.3). The total enzyme mass capacity limitation was determined by counting the number of metabolically active reactions in a simulation (under the relevant conditions), and extrapolating a linear fit of the cumulative enzyme mass from measured data vs. the number of enzymes measured. This yielded capacity bounds in the range 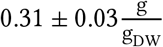 depending on the number of active reactions in the system. To estimate the robustness of this approach, we also used a coarser capacity bound of 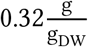 independent of the number of active reactions [52], but the results did not meaningfully change. Enzyme subunit stoichiometry was determined through Uniprot [53] and Complex Portal [54] annotations. Molar masses of proteins were taken from Uniprot. Turnover numbers were taken from [34], Michaelis constants from [15], and thermodynamic data from [1]. All simulations were performed using COBREXA.jl [55], a modern Julia Language [56] constraint-based analysis package. CPLEX [36] was used as the optimization solver. The models of Problems (P2), (P3), and (P4) were differentiated as described in the text. The specific model format is described below. Gradient descent was performed on Problem (P3) using an approximate, backtracking line search algorithm to find an appropriate step size. For Problem (P4), metabolite concentrations were estimated by minimizing the effect thermodynamics and saturation has on each enzyme, as described later. Visualizations were done using Makie [57].

### 5.2 Enzyme-constrained metabolic model formulation

Enzyme capacity and rate limitations represent a physiological constraint that shapes the resource allocation in a cell. Briefly, assuming that saturation and thermodynamic factors are negligible, the flux (*ν*) through an enzyme catalyzed reaction may be modeled by,

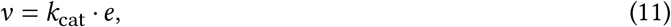

where *k*_cat_ is the turnover number of the associated enzyme, with concentration *e*. Multiple (related) algorithms incorporate this idea into constraint-based models [4, 6, 33]. Here, we focus on the GECKO formulation [6], but the results are generalizable to the other model variants.

For GECKO, the reactions of a constraint-based model are split into their forward and reverse components, so that all fluxes are positive. Additionally, each isozyme is also modeled as a distinct reaction. These unidirectional reactions are coupled in the optimization problem, so that the original reaction fluxes can be reconstructed from the newly created variables. By largely following the existing formulation, the resultant model is,

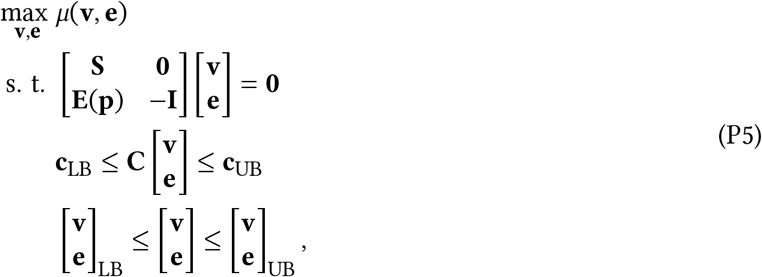

where the enzyme turnover numbers enter **E** as parameters (**p**), and the coupling matrix **C** allows the original bounds of the model to constrain the split reactions through **c**_LB_ and **c**_UB_. Additionally, enzyme capacity bounds can also be incorporated through **C**. Typically, the objective is some linear combination of fluxes (**v**) and enzyme concentrations (**e**), expressed through the function *µ*. Although each term in Equation (P5) could be a function of the parameters, for simplicity we assume that only **E** takes parameters.

For models that are purely enzyme-constrained, **E** is a sparse matrix, composed of terms,

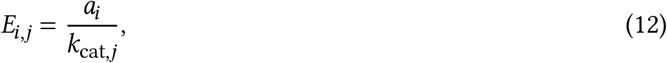

where the columns (*j*) correspond to enzyme-constrained reactions, and rows (*i*) are pseudo-protein mass balances (exactly like the GECKO formulation). Here *a*_*i*_ is the stoichiometric coefficient of the protein subunit catalyzing the associated reaction.

For models that incorporate thermodynamic and enzyme constraints, **E** is again a sparse matrix, but now composed of terms,

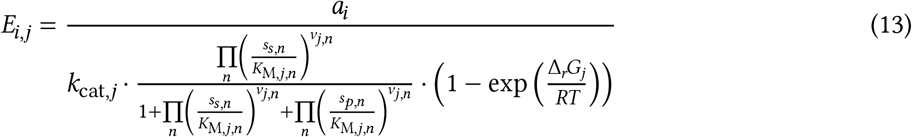

where the meaning of all the terms are the same as used throughout this work.

These problems can be converted into the standard form of Problem (P1), and then differentiated with respect to the associated parameters (most simply using symbolic or automatic differentiation).

### 5.3 Estimating intracellular concentrations

In Problem (P4), intracellular metabolite concentrations need to be supplied as parameters. In this work, we assumed that intracellular metabolite concentrations would be adjusted by the cell to ensure that enzymes work at maximum efficiency relative to saturation limitations. Briefly, the metabolite concentrations were estimated by minimizing the maximum impact of the saturation terms on enzyme velocity for all enzymes with Michaelis constant data, as shown in Problem (P6),

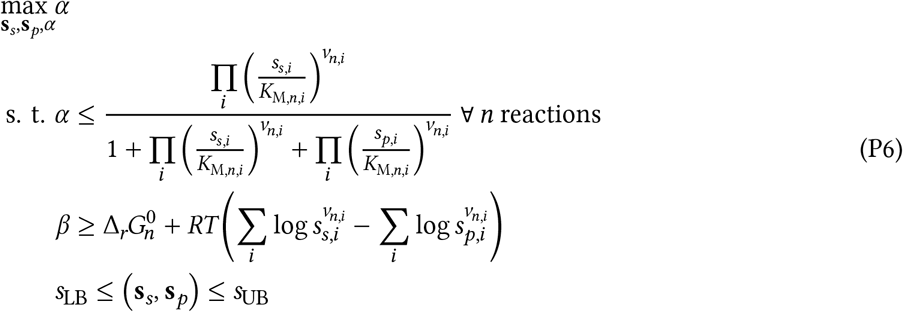

where *β* was set to 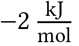 and represents a lower bound on the driving force of each reaction. Additionally, **s**_*s*_, **s**_*p*_ are the substrate and product concentrations relative to each reaction. All metabolite concentrations were bounded by *s*_LB_ = 10^−9^ M and *s*_UB_ = 0.1 M. This problem is nonlinear and nonconvex, a local solution was computed using KNITRO [58], using the metabolite concentrations from max-min driving force analysis [3] as starting points.

## 6 Acknowledgements

M. Besançon was partially supported through the Research Campus Modal funded by the German Federal Ministry of Education and Research (fund numbers 05M14ZAM, 05M20ZBM). This work was partially funded by the Deutsche Forschungsgemeinschaft (DFG, German Research Foundation) under Germany’s Excellence Strategy — EXC-2048/1 — project ID 390686111. This work was also partially supported by the European Union’s Horizon 2020 Programme under the PerMedCoE Project (www.permedcoe.eu) [951773].

## 7 Author Contributions

**St. Elmo Wilken**: Conceptualization, Data curation, Software, Methodology, Investigation, Writing - Original draft preparation. **Mathieu Besançon**: Methodology, Writing - review & editing. **Miroslav Kratochvíl**: Software, Writing - review & editing. **Chilperic Armel Foko Kuate**: Validation, Funding acquisition, Writing - review & editing. **Christophe Trefois**: Funding acquisition, Writing - review & editing. **Wei Gu**: Funding acquisition, Writing - review & editing. **Oliver Ebenhöh**: Validation, Funding acquisition, Writing - review & editing.

## References

[1] Avi Flamholz, Elad Noor, Arren Bar-Even, and Ron Milo. Equilibrator—the biochemical thermodynamics calculator. Nucleic acids research, 40(D1):D770–D775, 2012.

[2] Antje Chang, Lisa Jeske, Sandra Ulbrich, Julia Hofmann, Julia Koblitz, Ida Schomburg, Meina Neumann-Schaal, Dieter Jahn, and Dietmar Schomburg. Brenda, the elixir core data resource in 2021: new developments and updates. Nucleic Acids Research, 49(D1):D498–D508, 2021.

[3] Elad Noor, Arren Bar-Even, Avi Flamholz, Ed Reznik, Wolfram Liebermeister, and Ron Milo. Pathway thermodynamics highlights kinetic obstacles in central metabolism. PLoS computational biology, 10(2):e1003483, 2014.

[4] Roi Adadi, Benjamin Volkmer, Ron Milo, Matthias Heinemann, and Tomer Shlomi. Prediction of microbial growth rate versus biomass yield by a metabolic network with kinetic parameters. PLoS computational biology, 8(7):e1002575, 2012.

[5] Christopher S Henry, Linda J Broadbelt, and Vassily Hatzimanikatis. Thermodynamics-based metabolic flux analysis. Biophysical journal, 92(5):1792–1805, 2007.

[6] Benjamın J Sánchez, Cheng Zhang, Avlant Nilsson, Petri-Jaan Lahtvee, Eduard J Kerkhoven, and Jens Nielsen. Improving the phenotype predictions of a yeast genome-scale metabolic model by incorporating enzymatic constraints. Molecular systems biology, 13(8):935, 2017.

[7] Jingru Zhou, Yingping Zhuang, and Jianye Xia. Integration of enzyme constraints in a genome-scale metabolic model of aspergillus niger improves phenotype predictions. Microbial Cell Factories, 20(1):1–16, 2021.

[8] St. Elmo Wilken, Victor Vera Frazão, Nima P Saadat, and Oliver Ebenhöh. The view of microbes as energy converters illustrates the trade-off between growth rate and yield. Biochemical Society Transactions, 49(4):1663–1674, 2021.

[9] Gang Li, Yating Hu, Jan Zrimec, Hao Luo, Hao Wang, Aleksej Zelezniak, Boyang Ji, and Jens Nielsen. Bayesian genome scale modelling identifies thermal determinants of yeast metabolism. Nature communications, 12(1):1–12, 2021.

[10] Qasim K Beg, Alexei Vazquez, Jason Ernst, Marcio A de Menezes, Ziv Bar-Joseph, A-L Barabási, and Zoltán N Oltvai. Intracellular crowding defines the mode and sequence of substrate uptake by escherichia coli and constrains its metabolic activity. Proceedings of the National Academy of Sciences, 104(31):12663–12668, 2007.

[11] Daan H De Groot, Julia Lischke, Riccardo Muolo, Robert Planqué, Frank J Bruggeman, and Bas Teusink. The common message of constraint-based optimization approaches: overflow metabolism is caused by two growth-limiting constraints. Cellular and Molecular Life Sciences, 77(3):441–453, 2020.

[12] Dan Davidi, Elad Noor, Wolfram Liebermeister, Arren Bar-Even, Avi Flamholz, Katja Tummler, Uri Barenholz, Miki Goldenfeld, Tomer Shlomi, and Ron Milo. Global characterization of in vivo enzyme catalytic rates and their correspondence to in vitro kcat measurements. Proceedings of the National Academy of Sciences, 113(12):3401–3406, 2016.

[13] Yu Chen and Jens Nielsen. In vitro turnover numbers do not reflect in vivo activities of yeast enzymes. Proceedings of the National Academy of Sciences, 118(32), 2021.

[14] David Heckmann, Colton J Lloyd, Nathan Mih, Yuanchi Ha, Daniel C Zielinski, Zachary B Haiman, Abdelmoneim Amer Desouki, Martin J Lercher, and Bernhard O Palsson. Machine learning applied to enzyme turnover numbers reveals protein structural correlates and improves metabolic models. Nature communications, 9(1):1–10, 2018.

[15] Alexander Kroll, Martin KM Engqvist, David Heckmann, and Martin J Lercher. Deep learning allows genome-scale prediction of michaelis constants from structural features. PLoS biology, 19(10):e3001402, 2021.

[16] Edward J O’brien, Joshua A Lerman, Roger L Chang, Daniel R Hyduke, and Bernhard Ø Palsson. Genomescale models of metabolism and gene expression extend and refine growth phenotype prediction. Molecular systems biology, 9(1):693, 2013.

[17] Anne Goelzer, Jan Muntel, Victor Chubukov, Matthieu Jules, Eric Prestel, Rolf Nölker, Mahendra Mariadassou, Stéphane Aymerich, Michael Hecker, Philippe Noirot, et al. Quantitative prediction of genome-wide resource allocation in bacteria. Metabolic engineering, 32:232–243, 2015.

[18] Reinhart Heinrich and Stefan Schuster. The regulation of cellular systems. Springer Science & Business Media, 2012.

[19] Avlant Nilsson and Jens Nielsen. Metabolic trade-offs in yeast are caused by f1f0-atp synthase. Scientific reports, 6(1):1–11, 2016.

[20] Sophia Tsouka, Meric Ataman, Tuure Hameri, Ljubisa Miskovic, and Vassily Hatzimanikatis. Constraint-based metabolic control analysis for rational strain engineering. Metabolic Engineering, 66:191–203, 2021.

[21] Iván Domenzain, Benjamın Sánchez, Mihail Anton, Eduard J Kerkhoven, Aarón Millán-Oropeza, Céline Henry, Verena Siewers, John P Morrissey, Nikolaus Sonnenschein, and Jens Nielsen. Reconstruction of a catalogue of genome-scale metabolic models with enzymatic constraints using gecko 2.0. BioRxiv, 2021.

[22] Vassily Hatzimanikatis and James E Bailey. Mca has more to say. Journal of theoretical Biology, 182(3):233–242, 1996.

[23] John Villadsen, Jens Nielsen, and Gunnar Lidén. Bioreaction engineering principles. Springer Science & Business Media, 2011.

[24] Jeffrey D Orth, Ines Thiele, and Bernhard Ø Palsson. What is flux balance analysis? Nature biotechnology, 28(3):245–248, 2010.

[25] Stephen Boyd, Stephen P Boyd, and Lieven Vandenberghe. Convex optimization. Cambridge university press, 2004.

[26] Stephen Gould, Basura Fernando, Anoop Cherian, Peter Anderson, Rodrigo Santa Cruz, and Edison Guo. On differentiating parameterized argmin and argmax problems with application to bi-level optimization. arXiv preprint 1607.05447, 2016.

[27] Brandon Amos and J Zico Kolter. Optnet: differentiable optimization as a layer in neural networks. In International Conference on Machine Learning, pages 136–145. PMLR, 2017.

[28] Mathieu Blondel, Quentin Berthet, Marco Cuturi, Roy Frostig, Stephan Hoyer, Felipe Llinares-López, Fabian Pedregosa, and Jean-Philippe Vert. Efficient and modular implicit differentiation. arXiv preprint 2105.15183, 2021.

[29] Akshay Sharma, Mathieu Besançon, Joaquim Dias Garcia, and Benoît Legat. Flexible Differentiable Optimization via Model Transformations, 2022. URL: https://arxiv.org/abs/2206.06135.

[30] William Moses and Valentin Churavy. Instead of rewriting foreign code for machine learning, automatically synthesize fast gradients. In H. Larochelle, M. Ranzato, R. Hadsell, M. F. Balcan, and H. Lin, editors, Advances in Neural Information Processing Systems, volume 33, pages 12472–12485. Curran Associates, Inc., 2020. URL: https://proceedings.neurips.cc/paper/2020/file/9332c513ef44b682e9347822c2e457ac-Paper.pdf.

[31] J. Revels, M. Lubin, and T. Papamarkou. Forward-mode automatic differentiation in Julia. 1607.07892 [cs.MS], 2016. URL: https://arxiv.org/abs/1607.07892.

[32] Shashi Gowda, Yingbo Ma, Alessandro Cheli, Maja Gwóźzdź, Viral B. Shah, Alan Edelman, and Christopher Rackauckas. High-performance symbolic-numerics via multiple dispatch. ACM Commun. Comput. Algebra, 55(3):92–96, January 2022. ISSN: 1932-2240. doi: 10.1145/3511528.3511535. URL: https://doi.org/10.1145/3511528.3511535.

[33] Pavlos Stephanos Bekiaris and Steffen Klamt. Automatic construction of metabolic models with enzyme constraints. BMC bioinformatics, 21(1):1–13, 2020.

[34] David Heckmann, Anaamika Campeau, Colton J Lloyd, Patrick V Phaneuf, Ying Hefner, Marvic Carrillo-Terrazas, Adam M Feist, David J Gonzalez, and Bernhard O Palsson. Kinetic profiling of metabolic specialists demonstrates stability and consistency of in vivo enzyme turnover numbers. Proceedings of the National Academy of Sciences, 117(37):23182–23190, 2020.

[35] Jonathan M Monk, Colton J Lloyd, Elizabeth Brunk, Nathan Mih, Anand Sastry, Zachary King, Rikiya Takeuchi, Wataru Nomura, Zhen Zhang, Hirotada Mori, et al. Iml1515, a knowledgebase that computes escherichia coli traits. Nature biotechnology, 35(10):904–908, 2017.

[36] IBM ILOG Cplex. V20.1.0: user’s manual for cplex. International Business Machines Corporation, 2021.

[37] Elad Noor, Avi Flamholz, Wolfram Liebermeister, Arren Bar-Even, and Ron Milo. A note on the kinetics of enzyme action: a decomposition that highlights thermodynamic effects. FEBS letters, 587(17):2772–2777, 2013.

[38] Bryson D Bennett, Jie Yuan, Elizabeth H Kimball, and Joshua D Rabinowitz. Absolute quantitation of intracellular metabolite concentrations by an isotope ratio-based approach. Nature protocols, 3(8):1299–1311, 2008.

[39] Stefano Donati, Michelle Kuntz, Vanessa Pahl, Niklas Farke, Dominik Beuter, Timo Glatter, Jose Vicente Gomes-Filho, Lennart Randau, Chun-Ying Wang, and Hannes Link. Multi-omics analysis of crispriknockdowns identifies mechanisms that buffer decreases of enzymes in e. coli metabolism. Cell Systems, 12(1):56–67, 2021.

[40] Elad Noor, Avi Flamholz, Arren Bar-Even, Dan Davidi, Ron Milo, and Wolfram Liebermeister. The protein cost of metabolic fluxes: prediction from enzymatic rate laws and cost minimization. PLoS computational biology, 12(11):e1005167, 2016.

[41] Charles J Foster, Lin Wang, Hoang V Dinh, Patrick F Suthers, and Costas D Maranas. Building kinetic models for metabolic engineering. Current Opinion in Biotechnology, 67:35–41, 2021.

[42] Jan Schellenberger and Bernhard Ø Palsson. Use of randomized sampling for analysis of metabolic networks. Journal of biological chemistry, 284(9):5457–5461, 2009.

[43] Neal Parikh and Stephen Boyd. Proximal algorithms. Foundations and Trends in optimization, 1(3):127–239, 2014.

[44] Daan H de Groot, Josephus Hulshof, Bas Teusink, Frank J Bruggeman, and Robert Planqué. Elementary growth modes provide a molecular description of cellular self-fabrication. PLoS computational biology, 16(1):e1007559, 2020.

[45] Hugo Dourado and Martin J Lercher. An analytical theory of balanced cellular growth. Nature communications, 11(1):1–14, 2020.

[46] Ulrike Wittig, Renate Kania, Martin Golebiewski, Maja Rey, Lei Shi, Lenneke Jong, Enkhjargal Algaa, Andreas Weidemann, Heidrun Sauer-Danzwith, Saqib Mir, et al. Sabio-rk—database for biochemical reaction kinetics. Nucleic acids research, 40(D1):D790–D796, 2012.

[47] Arren Bar-Even, Elad Noor, Yonatan Savir, Wolfram Liebermeister, Dan Davidi, Dan S Tawfik, and Ron Milo. The moderately efficient enzyme: evolutionary and physicochemical trends shaping enzyme parameters. Biochemistry, 50(21):4402–4410, 2011.

[48] Feiran Li, L. Yuan, Hongzhong Lu, Gang Li, Yu Chen, Martin KM Engqvist, Eduard J Kerkhoven, and Jens Nielsen. Deep learning-based kcat prediction enables improved enzyme-constrained model reconstruction. Nature Catalysis:1–11, 2022.

[49] Yu Chen and Jens Nielsen. Mathematical modeling of proteome constraints within metabolism. Current Opinion in Systems Biology, 25:50–56, 2021.

[50] Nobuyoshi Ishii, Kenji Nakahigashi, Tomoya Baba, Martin Robert, Tomoyoshi Soga, Akio Kanai, Takashi Hirasawa, Miki Naba, Kenta Hirai, Aminul Hoque, et al. Multiple high-throughput analyses monitor the response of e. coli to perturbations. Science, 316(5824):593–597, 2007.

[51] Junyoung O Park, Sara A Rubin, Yi-Fan Xu, Daniel Amador-Noguez, Jing Fan, Tomer Shlomi, and Joshua D Rabinowitz. Metabolite concentrations, fluxes and free energies imply efficient enzyme usage. Nature chemical biology, 12(7):482–489, 2016.

[52] Alexander Schmidt, Karl Kochanowski, Silke Vedelaar, Erik Ahrné, Benjamin Volkmer, Luciano Callipo, Kevin Knoops, Manuel Bauer, Ruedi Aebersold, and Matthias Heinemann. The quantitative and condition-dependent escherichia coli proteome. Nature biotechnology, 34(1):104–110, 2016.

[53] UniProt Consortium. Uniprot: a worldwide hub of protein knowledge. Nucleic acids research, 47(D1):D506–D515, 2019.

[54] Birgit HM Meldal, Livia Perfetto, Colin Combe, Tiago Lubiana, João Vitor Ferreira Cavalcante, Hema Bye-A-Jee, Andra Waagmeester, Noemi Del-Toro, Anjali Shrivastava, Elisabeth Barrera, et al. Complex portal 2022: new curation frontiers. Nucleic Acids Research, 50(D1):D578–D586, 2022.

[55] Miroslav Kratochvıl, Laurent Heirendt, St. Elmo Wilken, Taneli Pusa, Sylvain Arreckx, Alberto Noronha, Marvin van Aalst, Venkata P Satagopam, Oliver Ebenhöh, Reinhard Schneider, et al. Cobrexa. jl: constraint-based reconstruction and exascale analysis. Bioinformatics, 38(4):1171–1172, 2022.

[56] Jeff Bezanson, Alan Edelman, Stefan Karpinski, and Viral B Shah. Julia: a fresh approach to numerical computing. SIAM review, 59(1):65–98, 2017.

[57] Simon Danisch and Julius Krumbiegel. Makie.jl: flexible high-performance data visualization for julia. Journal of Open Source Software, 6(65):3349, 2021. doi: 10.21105/joss.03349. URL: https://doi.org/10.21105/joss.03349.

[58] Richard H Byrd, Jorge Nocedal, and Richard A Waltz. K nitro: an integrated package for nonlinear optimization. In Large-scale nonlinear optimization, pages 35–59. Springer, 2006.

