## Supplement for "Interrogating the effect of enzyme kinetics on metabolism using differentiable constraint-based models"

July 11, 2022

#### S1 Differentiability considerations

The mapping from parameters to solution of an optimization problem is not differentiable everywhere. When the optimization problem is not strongly convex, the solution, if one exists, is not necessarily unique. A typical case in linear optimization is dual degeneracy, when a whole face of the feasible polyhedron is optimal; arbitrary small perturbations of the constraint matrix or objective will change the optimal set. Differentiating the model with respect to parameters is piecewise smooth differentiable. Like in the case of automatic differentiation, the produced result is a heuristic quantity at non-differentiable point and depends on the algorithm itself and not only on the mathematical function it computes [1]. We can regularize the model to improve the reliability or consistency of the computed derivatives, for instance by adding a small diagonal quadratic regularizer to the objective function $f(\mathbf{x}) + \frac{\mu}{2}\|\mathbf{x}\|_2^2$, ensuring that the problem is at least $\mu$-strongly convex, making the optimum unique. An alternative approach specific to our problems, is to remove all the sources of metabolic degeneracy to ensure that the solution being differentiated is unique. Once the optimal basis is unique, the problem is smoothly differentiable [2, 3]. To achieve this, the model is simulated, and all inactive reactions, genes and metabolites are removed. Thereafter, the model is simulated again, and then differentiated as explained in the main text. This approach ensures that the basis of the optimal solution is unique. Once the optimization problem is differentiable, it can be used to solve bi-level problems by differentiating the optimality conditions [4].

#### S2 Supplementary figures and tables

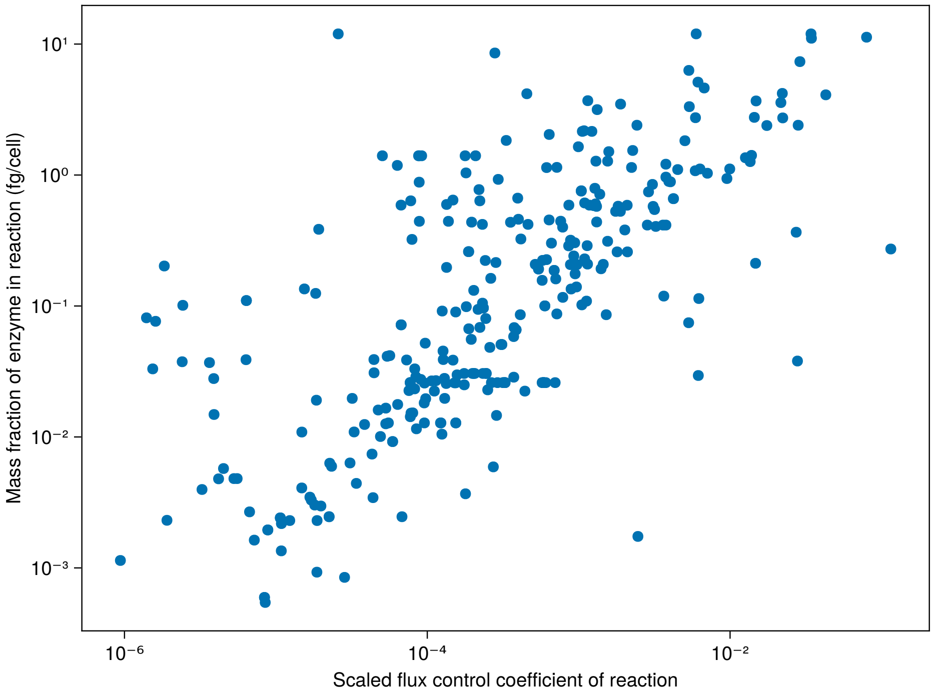


**Figure S1: Absolute proteomic quantification of enzyme mass per cell vs. flux control coefficients.** The direct scaled sensitivity of reaction $r$ with respect to its turnover number $k$ ($\frac{\partial \log r}{\partial \log k}$) is plotted against the absolute protein mass of $r$ for *E. coli* [5]. Transporter reactions are excluded from the analysis due to measurement bias effects. This sensitivities were determined using a glucose fed, aerobic, minimal media, enzyme constrained simulation of *E. coli* [6].

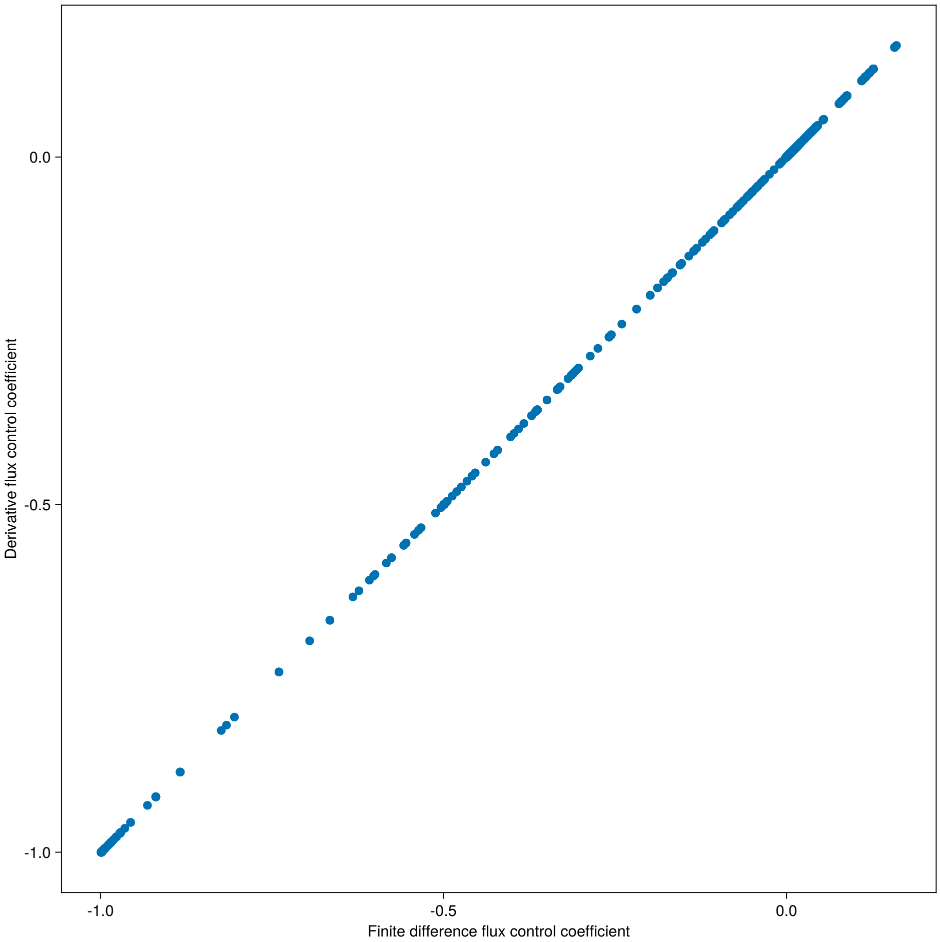


**Figure S2: Finite difference flux control coefficients vs. differential sensitivities.** Flux control coefficients can be calculated directly using the differentiation technique introduced in this work, or via a finite difference based approach [7]. In the latter case, the sensitivity can be iteratively calculated by perturbing a parameter, and inspecting the ratio of the perturbed solution to the reference (unperturbed) solution. Both techniques result in the exact same sensitivities, albeit the differentiation based approach is substantially more efficient, as only the reference optimization problem needs to be solved. This sensitivity comparison was done using a glucose fed, aerobic, minimal media, enzyme constrained simulation of *E. coli* [6].

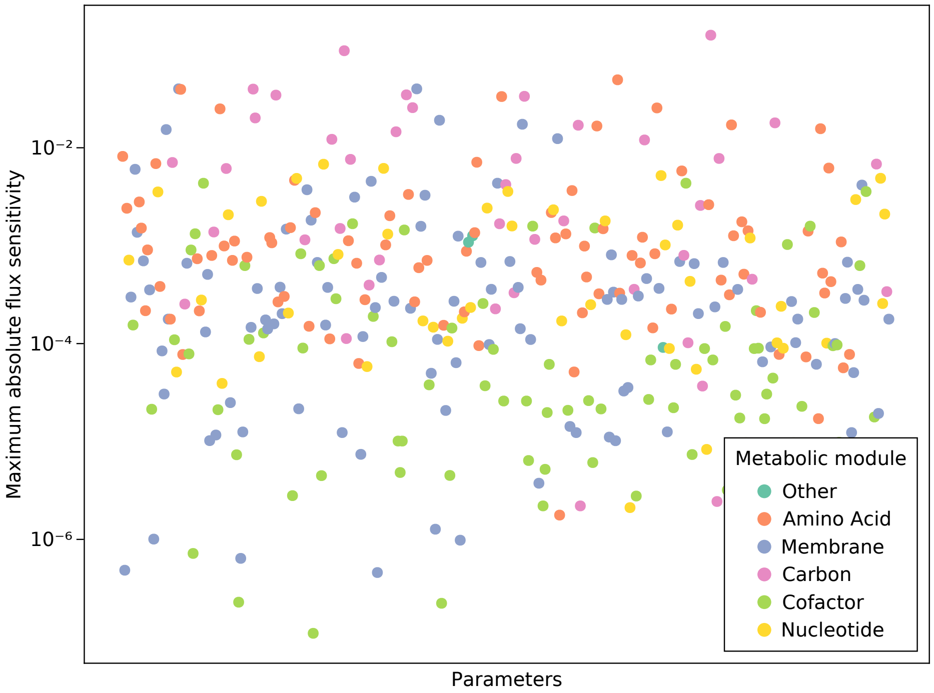


**Figure S3: The maximum scaled flux sensitivity of parameters is low for most parameters; metabolic fluxes are largely controlled by a few enzymes.** The maximum flux sensitivity of each parameter, colored by metabolic module of the parameter, over all reactions is shown for a glucose fed, aerobic, minimal media, enzyme constrained simulation of *E. coli* [6]. The maximum scaled flux sensitivity is small: only 27 parameters (out of 372) have a sensitivity on any reaction greater than $10^{-2}$.

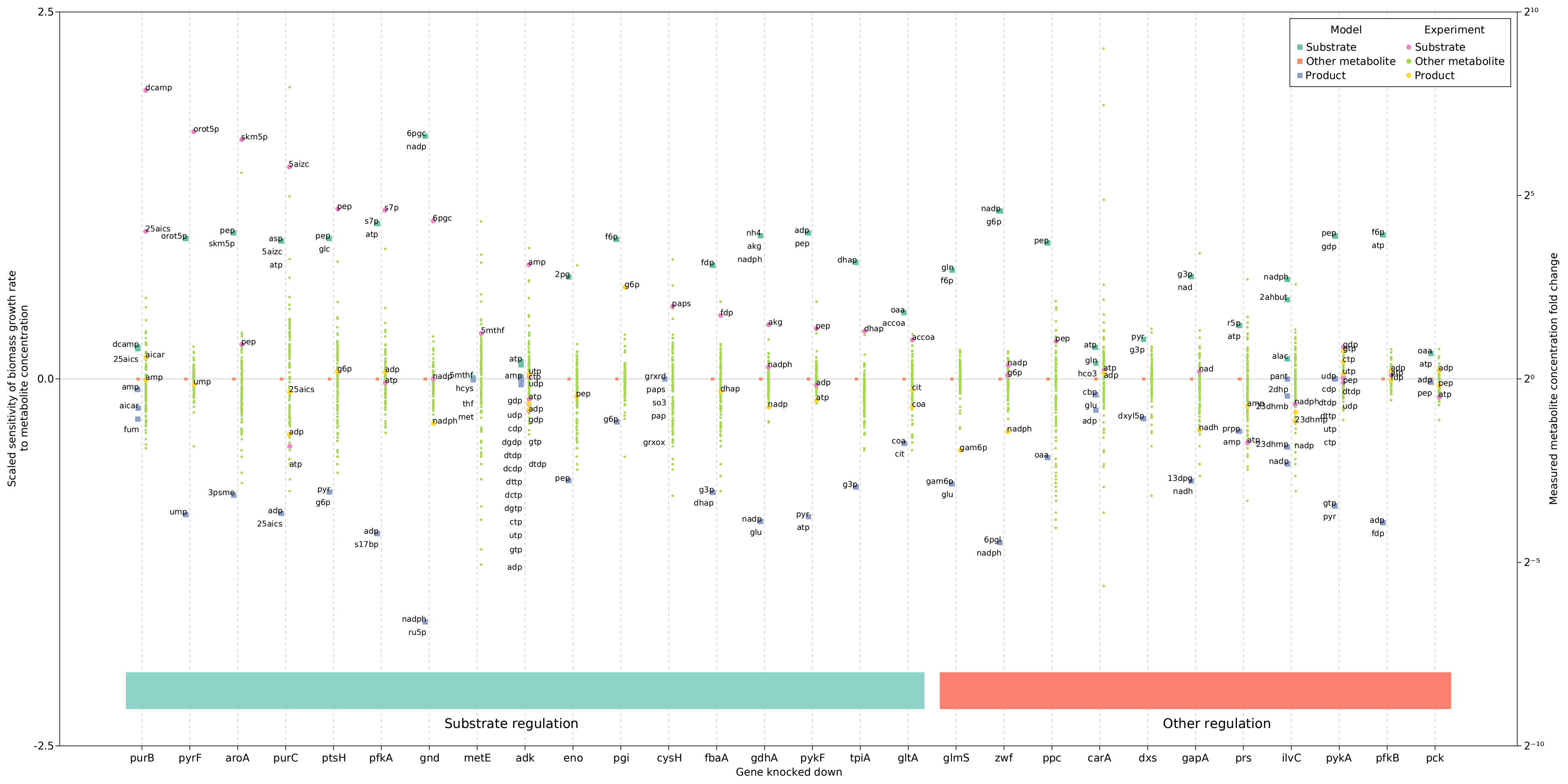


**Figure S4: Predicted biomass sensitivity to metabolites aligns with experimentally measured metabolome changes under genetic perturbations.** Each gene knockdown has two datasets associated with it. On the left y-axis, the simulated sensitivities of the biomass function to metabolite concentrations is shown after the associated gene is constrained to be five-fold less abundant than the wild type system. Substrate and product metabolite sensitivities are highlighted. On the right y-axis, experimental data of intracellular metabolite concentration fold changes observed after CRISPRi knockdowns of the associated genes are shown [8]. Substrate and product metabolites are also highlighted if measured. The gene abundance constraints experimentally observed match those simulated *in silico*. The groupings denoted by the horizontal blocks near the bottom of the figure separate the primary compensating mechanism observed *in vivo*. Briefly, "'Substrate regulation" denotes enzymes where the measured substrate metabolites played a large role in controlling the kinetics. "Other regulation" denotes cases where substrate concentration played a smaller role, e.g. allosteric regulation compensated for the knockdown, etc.

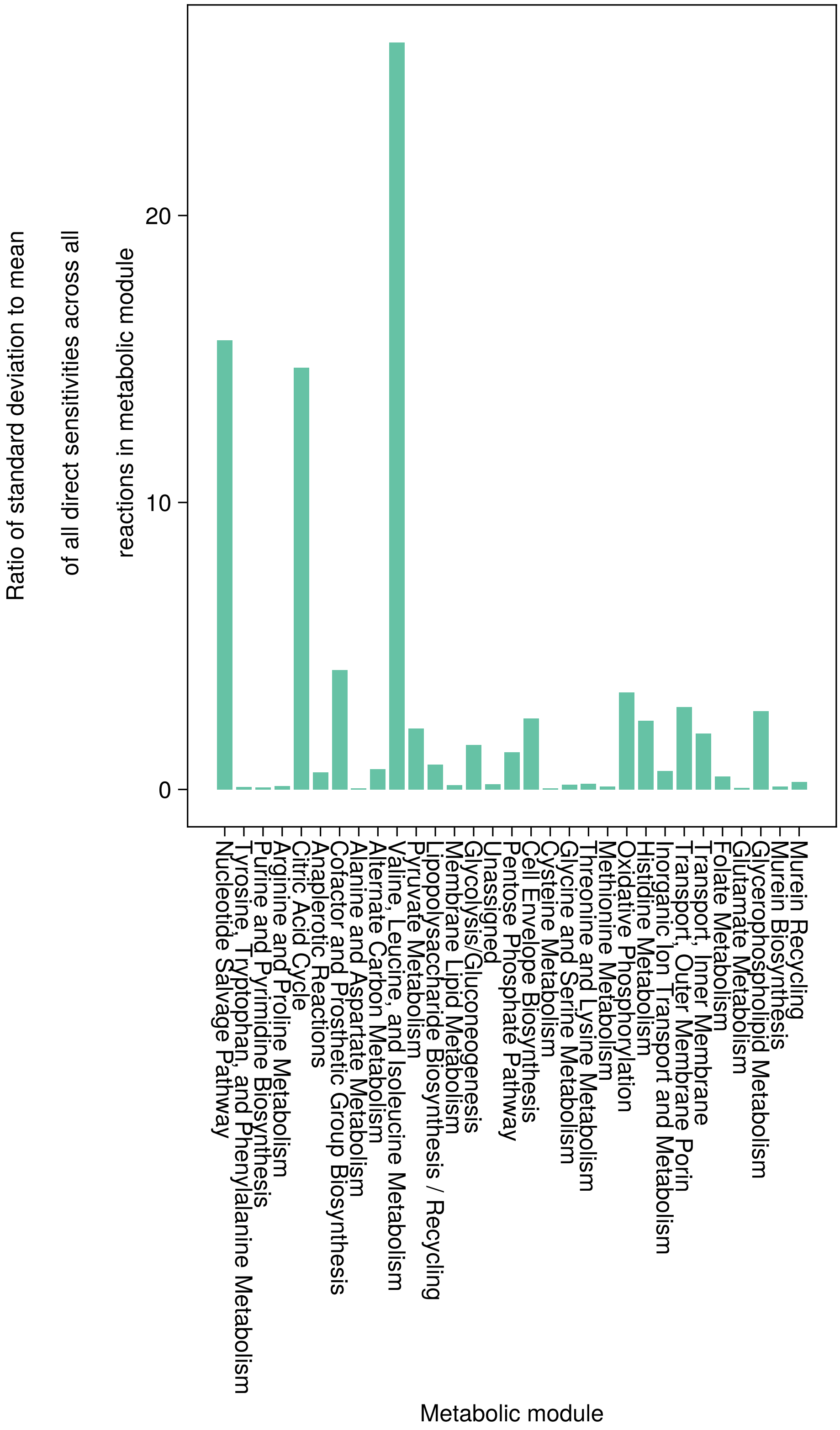


**Figure S5: Sensitivity analysis of different metabolic samples reveals that the sensitivities to parameters differ depending on the specific metabolic realization.** Flux variability analysis was performed on a glucose fed, aerobic, minimal media, enzyme constrained simulation of *E. coli* [6], where the biomass function was constrained to be within 10% of the optimum. Each sample collected represents an alternative metabolic state, yielding a relatively similar biomass growth prediction. The direct sensitivities (sensitivity of a reaction to its turnover number) were calculated, and the mean and standard deviation across all samples were found, and grouped by metabolic module of the reaction. The ratio of the standard deviation to mean for all these sensitivities are plotted, averaged and grouped by metabolic module. Most reaction groups show relatively little variability, but some prominent outliers show extreme variability in sensitivities under different metabolic realizations.

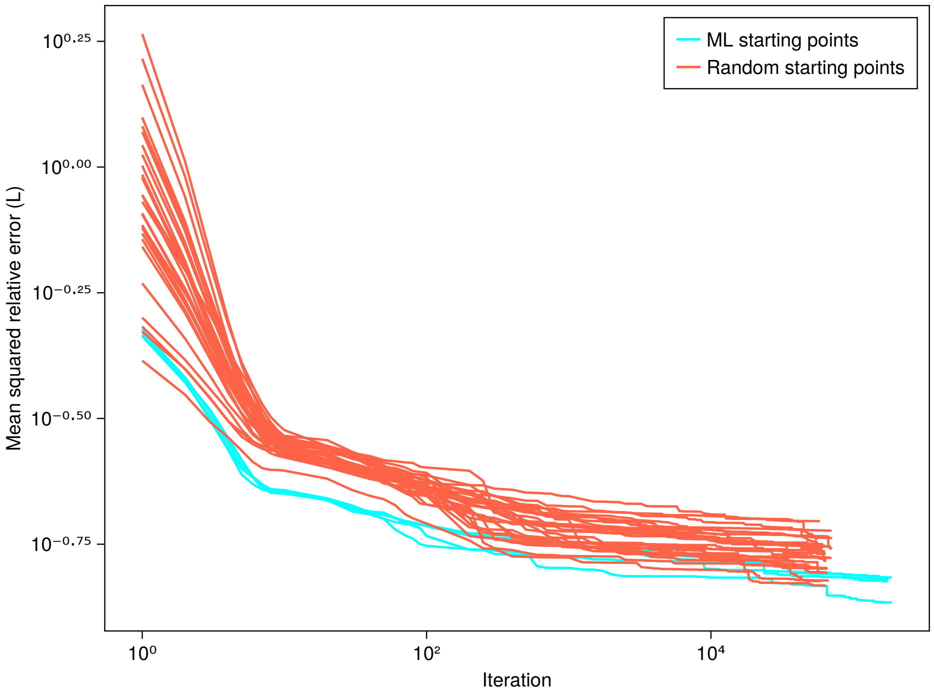


**Figure S6: Machine learning estimated turnover numbers are good starting points for gradient descent.** Gradient descent on the loss function, $L$, requires a starting estimate of the turnover numbers. Here, descent using wild type measurements from [9] are shown, with two classes of starting points. The first class are the machine learning (ML) estimates. The second class are turnover number starting values randomly sampled from a uniform distribution of each turnover number, with the lower/upper bound one order of magnitude lower/higher than the machine learning estimate. It is clear that the the machine learning estimates are good starting points, as their error is lower across almost all iterations.

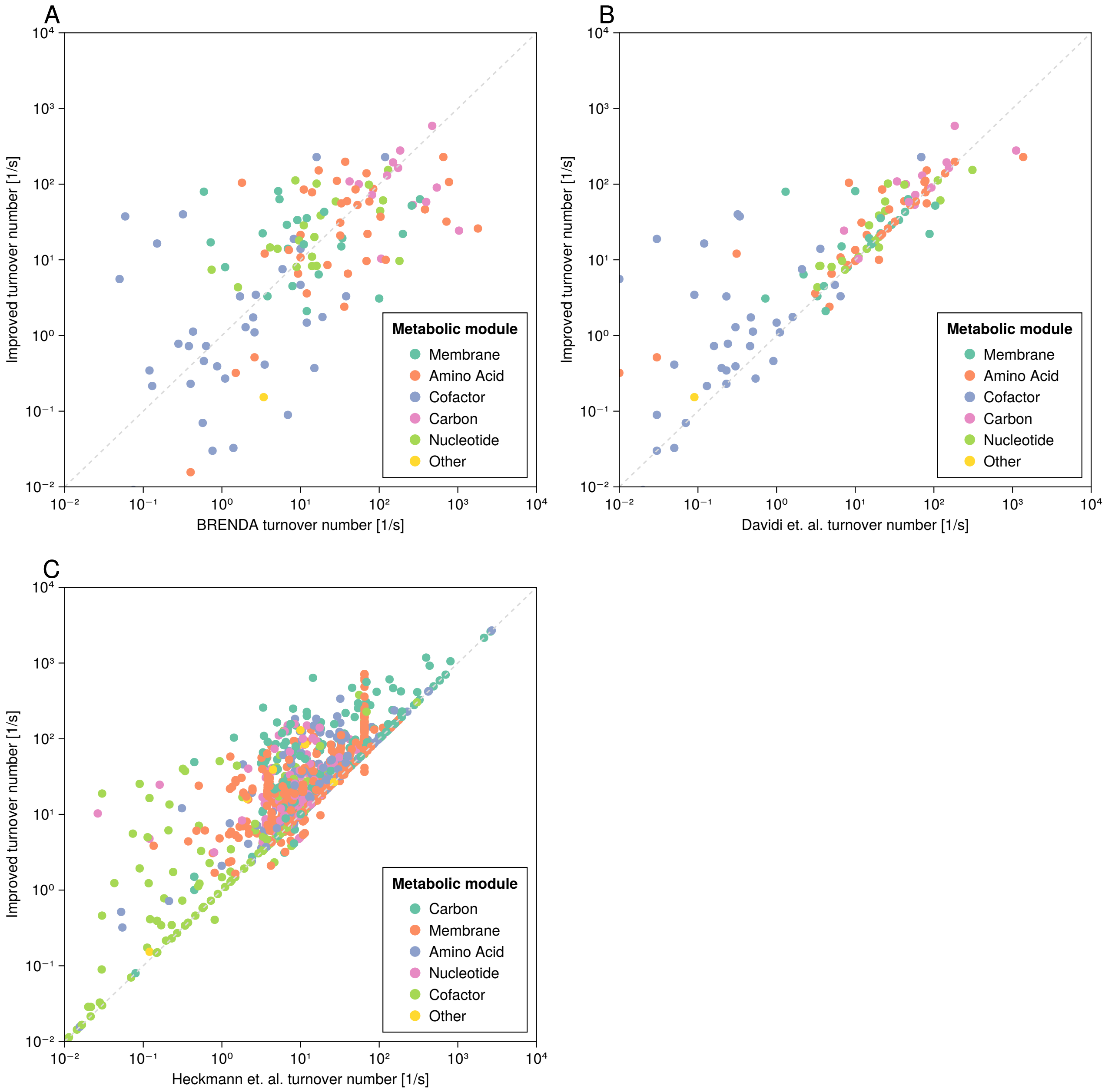


**Figure S7: Turnover number comparison between different sources.** Panel A reveals that there is no clear trend in how the gradient descent based approach changes the apparent turnover number relative to what is contained in BRENDA (BRENDA dataset was curated to remove high variability estimates) [10]. Similarly, Panel B shows no clear trend in how the turnover numbers were changed relative to those in Davidi et. al. (where an FBA-based approach was used to estimate apparent turnover numbers) [11]. Finally, Panel C shows that most apparent turnover numbers estimated by the gradient descent based approach were increased in magnitude relative to the machine learning based approach introduced by Heckmann et. al. [9].

### References


[1] Jerome Bolte and Edouard Pauwels. A mathematical model for automatic differentiation in machine learning. *Advances in Neural Information Processing Systems*, 33:10809–10819, 2020.

[2] Daniel De Wolf and Yves Smeers. Generalized derivatives of the optimal value of a linear program with respect to matrix coefficients. *European Journal of Operational Research*, 291(2):491–496, 2021.

[3] Robert M Freund. Postoptimal analysis of a linear program under simultaneous changes in matrix coefficients. In *Mathematical Programming Essays in Honor of George B. Dantzig Part I*, pages 1–13. Springer, 1985.
